# Absence of *MBP* 3’ UTR in Mice Disrupts mRNA Transport, Myelination, and Motor Learning

**DOI:** 10.64898/2025.12.23.696264

**Authors:** Joseph C. Nowacki, Haidyn L. Bulen, Lindsey M. Meservey, Hunter Richardson, Alex Valenzuela, Huy Nguyen, Zhuanfen Cheng, Hong Zeng, Meng-meng Fu

## Abstract

Myelin sheath maturation requires compaction, a cellular phenomenon mediated by local translation of myelin basic protein (MBP), which acts as a molecular zipper to join adjacent membranes and extrude the cytoplasm. Contrary to decades-old microinjection experiments indicating that *Mbp* mRNA transport is restricted to microtubules, we now show using smFISH that endogenous *Mbp* mRNA granules indeed localize along actin. To validate the *in vivo* necessity of *Mbp* mRNA transport and its dependence on the 3’ UTR (untranslated region), we replaced the endogenous *Mbp* 3’ UTR with polyA tail sequences. Though these mice have decreased *Mbp* mRNA levels, this alone could not account for their striking phenotypes – hypomyelination, baseline tremors, and motor learning defects. Thus, we cultured oligodendrocytes from these mice and found defects in both *Mbp* mRNA localization and local translation. These results demonstrate that the 3’ UTR of a locally translated structural protein is critical for both developmental and activity-induced myelination.

## INTRODUCTION

Efficient conduction of neuronal electrical signals along axons relies on myelin sheaths that insulate axons and define the Nodes of Ranvier. In order to produce myelin sheaths, oligodendrocytes in the central nervous system first extend many branched processes that contact axons (1). These processes then begin to ensheath and wrap axons in many concentric layers of myelin membrane (2). As myelin sheaths mature, they become compact, or devoid of cytoplasm (3). Compact myelin is composed of layer upon layer of adjacent membranes. Hence, in cross sections of axon bundles examined by electron microscopy (EM), compact myelin appears as dark, dense bands surrounding axons. Compaction is a unique and highly dynamic cellular transformation in which cytoplasm is extruded from the myelin sheath (4). The most prominent driver of the compaction process is a highly positively charged, small ∼20-kDa protein, MBP (myelin basic protein) (5), which is the second most abundant protein in the myelin sheath (6). Biophysical studies indicate that MBP joins adjacent layers within each wrap of the myelin sheath by acting as a molecular zipper, bringing together apposing membranes via hydrophobic interactions (4). Thus, MBP and myelin compaction are critically interdependent.

Indeed, *in vivo* studies demonstrate that MBP is essential for myelination. Classic studies in the *shiverer* mouse found that a large genomic deletion of Exons 3–7 in the *Mbp* gene (7, 8) results in a near complete loss of both *Mbp* mRNA and MBP protein expression (9), as well as failure to generate thick, compact myelin (10–12). As a result, homozygous *shiverer* mice exhibit tremors as early as postnatal days P10–15, seizures from P30 onward, and mortality between P50–100 (13, 14). However, heterozygous mice do not display any behavioral abnormalities even though they express only about half the amount of MBP protein as wildtype animals (9, 14–17).

Consistent with MBP’s role in joining apposed membranes (4), the site of MBP production is heavily regulated through mRNA transport and local translation. Expression of MBP in the cell body can be deleterious, aberrantly driving adhesion between the plasma membrane with organelles like the endoplasmic reticulum (4). Thus, local translation of MBP that is restricted to distal regions of oligodendrocytes limits compaction to the myelin sheath. Historically, MBP was one of the first proteins discovered to be locally translated in purified myelin fractions devoid of cell bodies or nuclei (18). Later, microinjection of fluorescent mRNA demonstrated that *Mbp* mRNA granules are transported away from the cell body and into cellular processes along microtubules (19–21). Microtubules are found both along the proximal branched processes and inside the myelin sheath (22–24). Further, they are essential for organelle (25) and mRNA transport, and disruptions in their organization or local nucleation result in thinner, shorter myelin sheaths (26). The anterograde transport of *Mbp* mRNA relies on both kinesin KIF1B (27) and dynein/dynactin motors, which engage in bidirectional transport, moving in a back-and-forth manner to distribute *Mbp* mRNA throughout the oligodendrocyte (28).

Importantly, it is unclear whether the *Mbp* 3’ UTR is necessary for its mRNA transport and local translation in an *in vivo* mammalian system. Indeed, classic rescue experiments in *shiverer* mice challenge this idea, because *Mbp* CDS (coding DNA sequence) transcripts devoid of 3’ UTR are sufficient to rescue the *shiverer* phenotype (29, 30). To test this, we targeted the 3’ untranslated region (3’ UTR) of *Mbp*, a region that is necessary for *Mbp* mRNA transport in cultured oligodendrocytes (31). An effective strategy to generate transgenic mice defective for mRNA transport is by replacing the 3’ UTR region with a generic polyA sequence, such as SV40 (simian virus) polyA. The SV40 polyA is commonly used in plasmids like pEGFP for the expression of tagged fluorescent proteins. This transgenic approach has been used previously to study *importin β1* and *mTOR* local translation in the context of axonal injury (Perry et al., 2012; Terenzio et al., 2018). Importantly, since the CDS is left intact, neither mouse model resulted in changes in levels of mRNA expression. Knocking out the *Importin β1* mRNA 3’ UTR reduces this mRNA’s localization to axons but also reduces the transcriptional response to axonal injury and delays recovery of motor function after injury *in vivo* (32). Similarly, the 3’ UTR of *mTOR* is also necessary for *mTOR* mRNA localization and local translation in injured axons (33). Thus, we took a similar approach of replacing endogenous *Mbp* 3’ UTR with SV40 polyA and generated a mouse that we call the *Mbp* 3’ UTR “knockout” (KO).

Here, we demonstrate that *Mbp* 3’ UTR KOs have aberrant localization of *Mbp* mRNA and lower levels of MBP protein. This is evident in both traditional 2D (two-dimensional) flat oligodendrocyte cultures as well as in 3D (three-dimensional) cultures that use microfibers to mimic the geometry of axons and generate nascent myelin sheaths. Strikingly, *Mbp* 3’ UTR KOs exhibit constant basal tremors. Furthermore, in a series of behavioral assays, we found that *Mbp* 3’UTR KOs also have gait defects, locomotor coordination defects, and motor learning deficits. Together, these data indicate that lack of the 3’ UTR of an abundant structural protein results in critical changes in development and basal neurological function.

## RESULTS

### *Mbp* mRNA localizes along actin

Classic studies with microinjected fluorescent mRNA indicated that *Mbp* mRNA transport occurs solely along microtubules in an actin-independent manner (20). To revisit this observation using modern approaches, we combined smFISH (single molecule fluorescent *in situ* hybridization) with spinning-disk confocal microscopy. We performed these experiments in primary oligodendrocytes isolated from P5 rat brains via the immunopanning method and differentiated them across a 6-day period. To observe changes in oligodendrocyte maturation, we combined *Mbp* mRNA smFISH with immunostaining against tubulin and fluorescent phalloidin labeling to visualize F-actin. On DIV 1 (days *in vitro* 1) and DIV 2, *Mbp* mRNA signal was indiscernible from the cytoplasmic background and we observed no *Mbp* mRNA-positive puncta (Fig. S1A). On DIV 3, a detectable *Mbp* mRNA smFISH signal appeared, concentrated in the cell body and proximal processes (Fig. S1A). By DIV 4, smFISH strikingly showed that endogenous *Mbp* mRNA granules are distributed as puncta along both microtubules and actin (Fig 1A). Thus, in addition to their transport along microtubules, *Mbp* mRNA granules also rely on actin for their localization.

**Figure 1.**
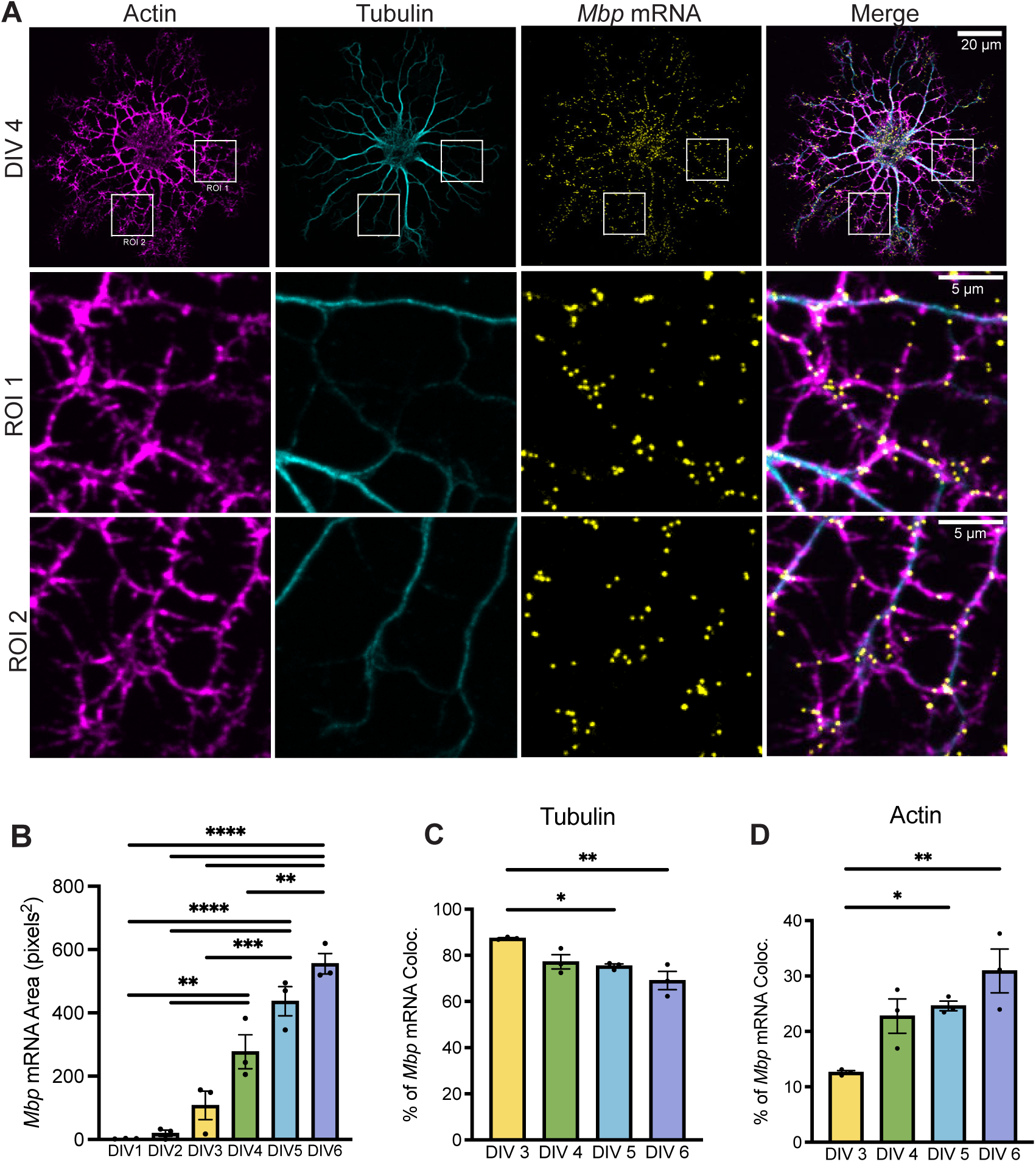
*Mbp* mRNA is transported along actin. (A) Confocal micrographs (60X) of DIV 4 immunopanned rat primary oligodendrocytes immunostained against tubulin, stained for actin using phalloidin, and smFISH against *Mbp* mRNA. *Mbp* mRNA colocalizes with tubulin and phalloidin immunopositive cytoskeleton. (B) *Mbp* mRNA area from DIV 1 to DIV 6 (One-way ANOVA with Tukey’s multiple comparisons test, n=3 individual experiments, 12–21 cells per DIV). (C) Percent of *Mbp* mRNA colocalized with tubulin signal from DIV 3–6 (P*=0.043, P**=0.004, One-way ANOVA with Tukey’s multiple comparisons test, n=3 experiments, 11–15 cells per DIV). (D) Percent of *Mbp* mRNA colocalized with phalloidin signal from DIV 3–6 (P*=0.043, P**=0.004, One-way ANOVA with Tukey’s multiple comparisons test, n=3 experiments, 11–15 cells per DIV). Data represents mean ± SEM. *P < 0.05, **P < 0.01, ***P < 0.001, ****P< 0.0001.

To gain insight into the temporal relationship between *Mbp* mRNA distribution along microtubules vs. actin, we quantified smFISH images of oligodendrocytes as they matured from DIV 1–6. As oligodendrocytes differentiation progressed, *Mbp* mRNA expression levels and localization followed a stereotyped pattern. From DIV 1 to DIV 6, the surface area occupied by *Mbp* smFISH puncta increased, indicative of an increase in number of *Mbp* mRNA granules (Fig. 1B, S1A). Compared to DIV 3 oligodendrocytes, DIV 6 oligodendrocytes had increased *Mbp* mRNA localization in distal regions >20 μm from the cell body (Fig. S1B). From DIV 3 to DIV 6, as more *Mbp* mRNA granules accumulate, their colocalization with microtubules decreases (Fig. 1C) while colocalization with actin increases (Fig. 1D), indicating that *Mbp* mRNA granules are likely first transported along microtubules, then along actin. Taken together, our data support a stereotyped pattern of oligodendrocyte development, in which an increasing number of *Mbp* granules are produced and distributed distally along microtubules and actin as these cells mature.

### MBP translation is temporally regulated

To determine how *Mbp* mRNA distribution relates to the timing of MBP protein translation, we simultaneously visualized endogenous *Mbp* mRNA using smFISH and MBP protein using antibody staining. Starting at DIV 4, MBP staining outside of the cell body can be observed in some distal areas of the cell (Fig. 2A and 2B). From DIV 4 to DIV 6, MBP protein area increased significantly each day (Fig. 2B and 2C). Compared to DIV-4 oligodendrocytes, DIV-6 oligodendrocytes had elevated MBP protein signal in distal regions >25 μm from the cell body (Fig. 2C). Together, our *Mbp* mRNA and MBP protein localization data reveal a coordinated sequence of events, starting with elevated transcription, transport along microtubules and actin, and ending with local translation of MBP.

**Figure 2.**
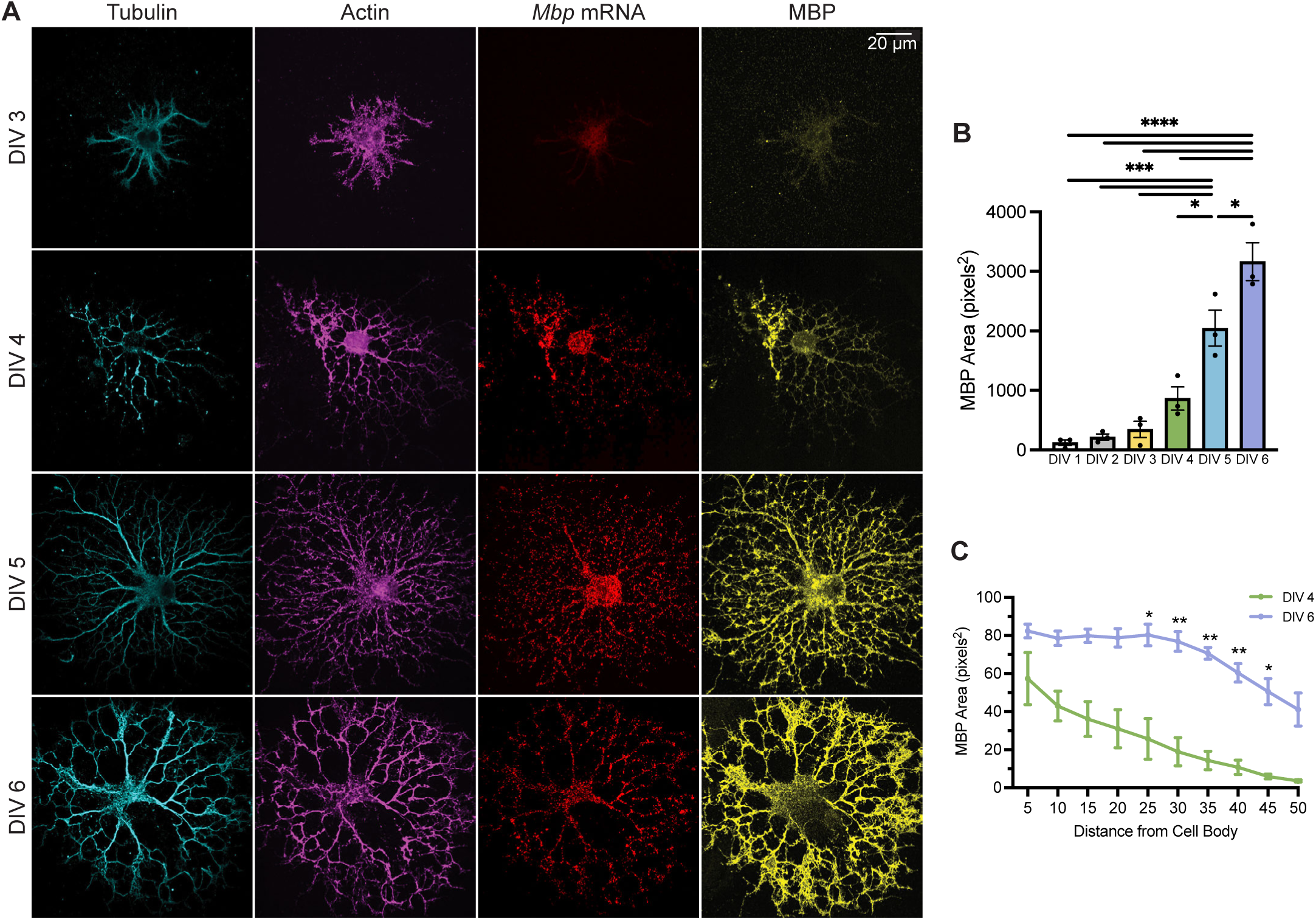
MBP translation is temporally regulated. (A) Confocal micrographs (60X) of DIV 3, 4, 5, 6 immunopanned rat primary oligodendrocytes immunostained against tubulin and MBP and stained for actin using phalloidin. (B) MBP protein area from DIV 1 to DIV 6 (One-way ANOVA with Tukey’s multiple comparisons test, n=3 individual experiments, 14–19 cells per DIV). (C) MBP protein area in concentric rings around the cell nucleus (Two-way ANOVA with Šídák’s multiple comparisons test, n=3 individual experiments, 14–17 cells per DIV). Data represents mean ± SEM. *P < 0.05, **P < 0.01, ***P < 0.001, ****P< 0.0001.

### A transgenic mouse lacking the endogenous *Mbp* 3’ UTR

Since both *Mbp* mRNA transport and translation depend on its 3’ UTR (31, 34), we next asked whether we could test this dependence *in vivo*. To do this, we generated a transgenic mouse by replacing the endogenous 3’ UTR with a stabilizing SV40 polyA sequence (Fig. 3A), which is commonly used for stable expression of fluorescently tagged proteins via mammalian vectors such as pEGFP. Although this strategy involves knocking in the SV40 polyA cassette, it effectively constitutes a 3’ UTR loss-of-function; therefore, we refer to this line as the *Mbp* 3’ UTR “knockout” (KO) model.

**Figure 3.**
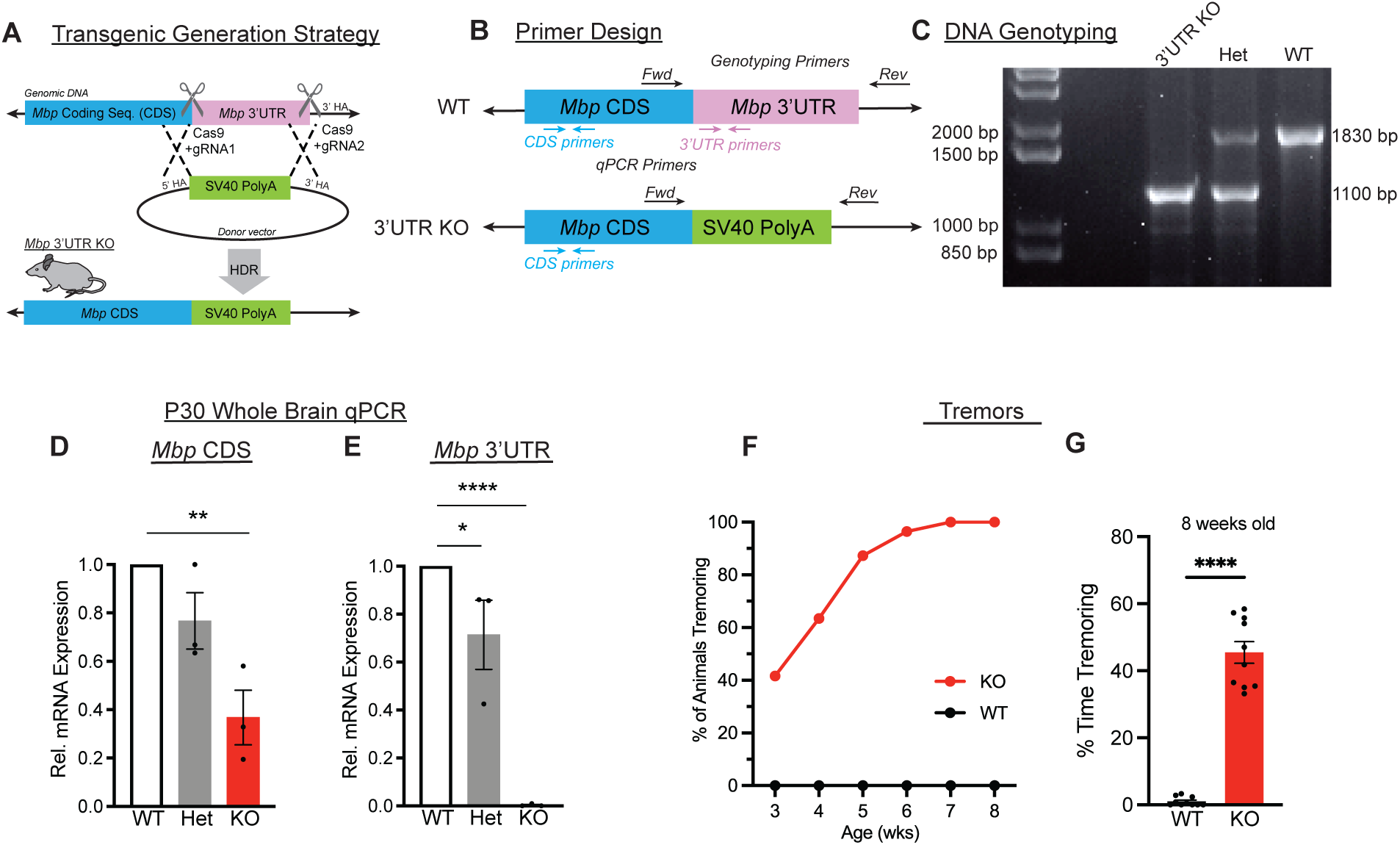
A transgenic mouse lacking the endogenous *Mbp* 3’UTR. (A) Transgenic generation strategy diagram. Guide RNAs were generated for the region immediately surrounding the *Mbp* 3’ UTR (3’ untranslated region). A donor construct containing 4 SV40 PolyA sequences with flanking regions that are homologous to the region flanking the *Mbp* 3’ UTR was generated. The *Mbp* 3’ UTR “knockout” was generated after homology-directed repair (HDR) or homologous recombination. (B) Primer design for DNA genotyping of wildtype and *Mbp* 3’ UTR KO. (C) DNA gel of wildtype, heterozygous, and *Mbp* 3’ UTR KO mice. (D) Relative *Mbp* mRNA CDS expression from P30 mice (WT vs. Het P=0.766, WT vs. KO P=0.0022, Het vs. KO P=0.0329, One-way ANOVA with Tukey’s multiple comparisons test, n=3 animals per genotype). (E) Relative *Mbp* 3’ UTR expression from P30 mice (WT vs. Het P=0.0606, WT vs. KO P < 0.0001, Het vs. KO P=0.0008, Kruskal Wallis test). (F) Tremor onset in *Mbp* 3’ UTR KO mice begins postnatally. Age vs. % of animals tremoring (n=9–11 animals per genotype). (G) Percentage time tremoring (P<0.0001, Mann-Whitney test, n=10 animals per genotype). Data represents mean ± SEM. *P < 0.05, **P < 0.01, ***P < 0.001, ****P< 0.0001.

To confirm the correct generation of this mouse, we sequenced the *Mbp* genomic region and developed a standardized PCR (polymerase chain reaction) genotyping protocol. We designed primers flanking the endogenous *Mbp* 3’ UTR region as well as the knocked in SV40 region (Fig. 3B), which had band sizes ∼1.8 kb and ∼1.1 kb, respectively. As expected, the larger 3’ UTR band is only present in wildtype mice while only the smaller SV40 band is present in homozygous *Mbp* 3’ UTR KO mice (Fig. 3C).

Next, to ascertain the effect of replacing the 3’ UTR on *Mbp* mRNA expression levels, we performed quantitative PCR (qPCR) using validated primer pairs targeting either the coding sequence (CDS) or the 3’ UTR (Fig. S2F–I). As expected, *Mbp* 3’ UTR transcripts were undetectable in homozygous mice (Fig. 3E, S2D, and S2E). In heterozygous *Mbp* 3’ UTR knockout mice, *Mbp* CDS mRNA is expressed at approximately 80% of wildtype levels (Fig. 3D). This perhaps suggests compensation in heterozygous *Mbp* 3’ UTR KO mice via upregulation of *Mbp* transcription.

Looking at homozygous *Mbp* 3’ UTR KO mice whole brains, we were surprised to find that they had less *Mbp* CDS expression, at both P14 and P30, to approximately 40% of wildtype levels (Fig. 3D and Fig. S2A). Region specific analyses from P30 brains further confirmed decreased *Mbp* CDS expression in the corpus callosum, a major white-matter tract (Fig. S2B), and in the spinal cord (Fig. S2C). This reduction contrasts with *importin β1* and *mTOR* 3’ UTR KO mice, which were generated using similar strategies but exhibited no change in CDS mRNA levels (32, 33). Thus, substitution of the *Mbp* 3’ UTR with SV40 polyA disrupts normal *Mbp* mRNA expression, suggesting that the endogenous 3’ UTR contributes to maintaining physiological *Mbp* mRNA levels.

Though *Mbp* 3’ UTR KO mice have reduced *Mbp* CDS mRNA expression, a reduction in *Mbp* transcript alone does not necessarily produce a behavioral phenotype. Importantly, a recent study that manipulated *Mbp* enhancer regions yielded two mouse lines with ∼50% reduction in *Mbp* mRNA; these mice had no alterations in their 3’ UTR and no discernible phenotypes (35). In addition, *shiverer* heterozygotes also express less *Mbp* mRNA (36) and are generally described as behaviorally normal with no overt clinical phenotype (15–17). However, we were surprised to find in routine handling of homozygous *Mbp* 3’ UTR KOs that they are visually distinct from wildtype and heterozygous mice. As early as 3 weeks of age, ∼40% of homozygous *Mbp* 3’UTR KO mice display prominent basal tremors. By 7–8 weeks of age, all homozygous *Mbp* 3’ UTR KO mice exhibit tremors (Fig. 3F), both at rest and while walking (Fig. 3G). Thus, the decrease in *Mbp* CDS mRNA alone (Fig. 3D) is insufficient to explain this severe tremor phenotype, implying an additional functional disruption such as perturbed *Mbp* mRNA localization or local translation.

### *Mbp* 3’ UTR KO oligodendrocytes have *Mbp* mRNA transport defects

Next, we asked whether absence of the 3’ UTR results in changes in *Mbp* mRNA transport or local translation. Given that myelination *in vivo* occurs in 3D, we generated 3D primary oligodendrocyte cultures using electrospun microfibers with 2-μm diameters that mimic the shape of axons. Oligodendrocytes grown for 2 weeks (DIV 14) extend processes that ensheath then wrap around the microfibers, forming nascent myelin sheaths (37). We visualized these 3D oligodendrocyte cultures via tubulin and MBP immunostaining as well as *Mbp* mRNA smFISH (Fig. 4A and 4B). Quantification of microtubule length revealed no significant difference between wildtype and homozygous *Mbp* 3’ UTR KO oligodendrocytes (Fig. 4C), suggesting comparable sheath lengths. However, we found dramatically reduced distribution of *Mbp* mRNA (Fig. 4D) and MBP protein (Fig. 4E) along the nascent myelin sheaths. Thus, these 3D culture experiments indicate that although the microtubules along nascent myelin sheaths are intact, the lack of the 3’ UTR impairs *Mbp* mRNA transport and results in downstream deficits in MBP local translation.

**Figure 4.**
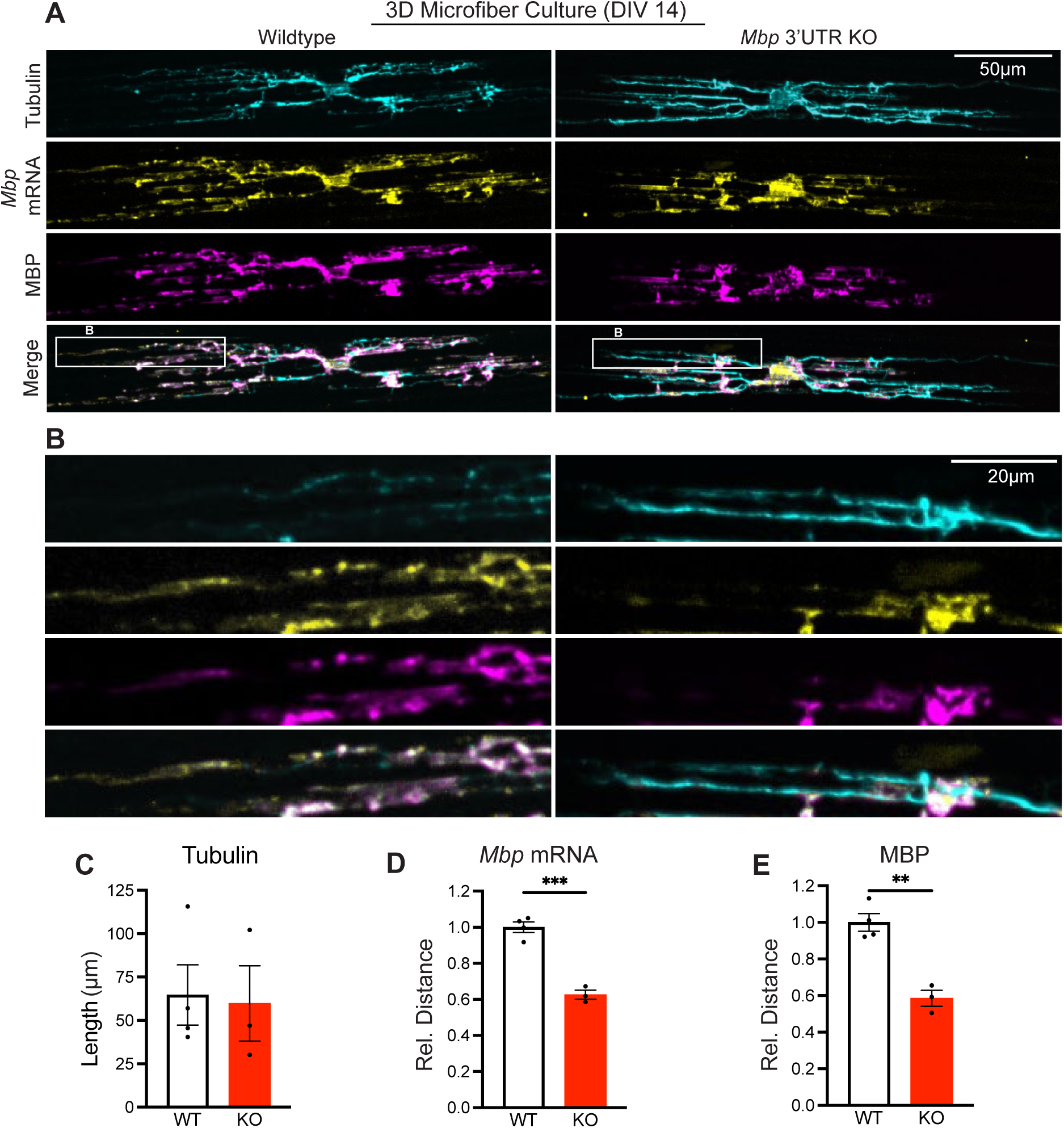
*Mbp* 3’ UTR KO oligodendrocytes show deficits in *Mbp* mRNA transport and local translation. (A) Max projection of confocal micrographs (20x) of DIV14 primary oligodendrocytes cultured from WT or *Mbp* 3’ UTR KO mice on 3D microfibers. Cells were stained against tubulin, and MBP, and smFISH was performed for *Mbp* mRNA. Merged images show all channels (n=3 separate experiments, 3–11 cells per genotype). (B) Box region from (A) zoomed in on individual processes of primary oligodendrocytes cultured from wildtype and *Mbp* 3’ UTR KO mice, respectively. (C) Tubulin length (P=0.86, Student’s t-test). (D) Relative distance of *Mbp* mRNA along tubulin cytoskeleton (P=0.0003, Student’s t-test). (E) Relative distance of MBP along tubulin cytoskeleton (P=0.0017, Student’s t-test). Data represents mean ± SEM. *P < 0.05, **P < 0.01, ***P < 0.001, ****P< 0.0001.

Nevertheless, we were surprised that *Mbp* mRNA lacking the 3’ UTR moved along the myelin sheath at all. Thus, to investigate how 3’ UTR-less *Mbp* mRNA translocated along the myelin sheath, we cultured flat 2D oligodendrocytes, which allow us to image with higher resolution than 3D cultures. These cells were differentiated for 4 days, then *Mbp* mRNA CDS was visualized using smFISH (Fig. S3A). We were intrigued to find large, round *Mbp* mRNA structures with diameters of 1–2 μm. We asked whether these are lysosomes, which have been demonstrated to associate with hitchhiking mRNAs in neurons (38, 39). Thus, we co-stained against the lysosomal marker LAMP1 and confirmed that *Mbp* CDS mRNA and LAMP1 indeed colocalized (Fig. S3A). This unexpected result suggests that in the absence of its endogenous 3’ UTR, *Mbp* mRNA transcripts containing the CDS and SV40 polyA may adopt other non-canonical compensatory transport mechanisms.

### *Mbp* 3’ UTR KO mice are hypomyelinated

We next examined MBP protein expression in intact brain tissue to assess whether myelination was altered at the cellular or regional level. As early as P21, decreased MBP staining was observed in homozygous *Mbp* 3’ UTR KO brains (Fig. S4A). High magnification images of the cerebellum also showed prominent lack of elongated myelin sheath structures (Fig. S4B). At P90, we found that homozygous mice have decreased MBP staining in the hippocampus, cerebellum, and cortex (Fig. S4C, 5A, and 5B). This pattern is particularly pronounced in the cerebellum, where sheath area was significantly reduced throughout the homozygous *Mbp* 3’ UTR KO cerebellum (Fig. 5C), as was the thickness of the long fingerlike projections of white matter that extend into the cerebellum (Fig. 5D). Using high-magnification (60X) images to further investigate sheath-level changes in myelination (Fig. 5E), we found that myelin sheath MBP intensity (Fig. 5F) and MBP area (Fig. 5G) is significantly decreased in homozygous *Mbp* 3’ UTR KO mice. Together, these findings demonstrate that *Mbp* 3’ UTR is necessary for myelin sheath formation and maintenance *in vivo*.

**Figure 5.**
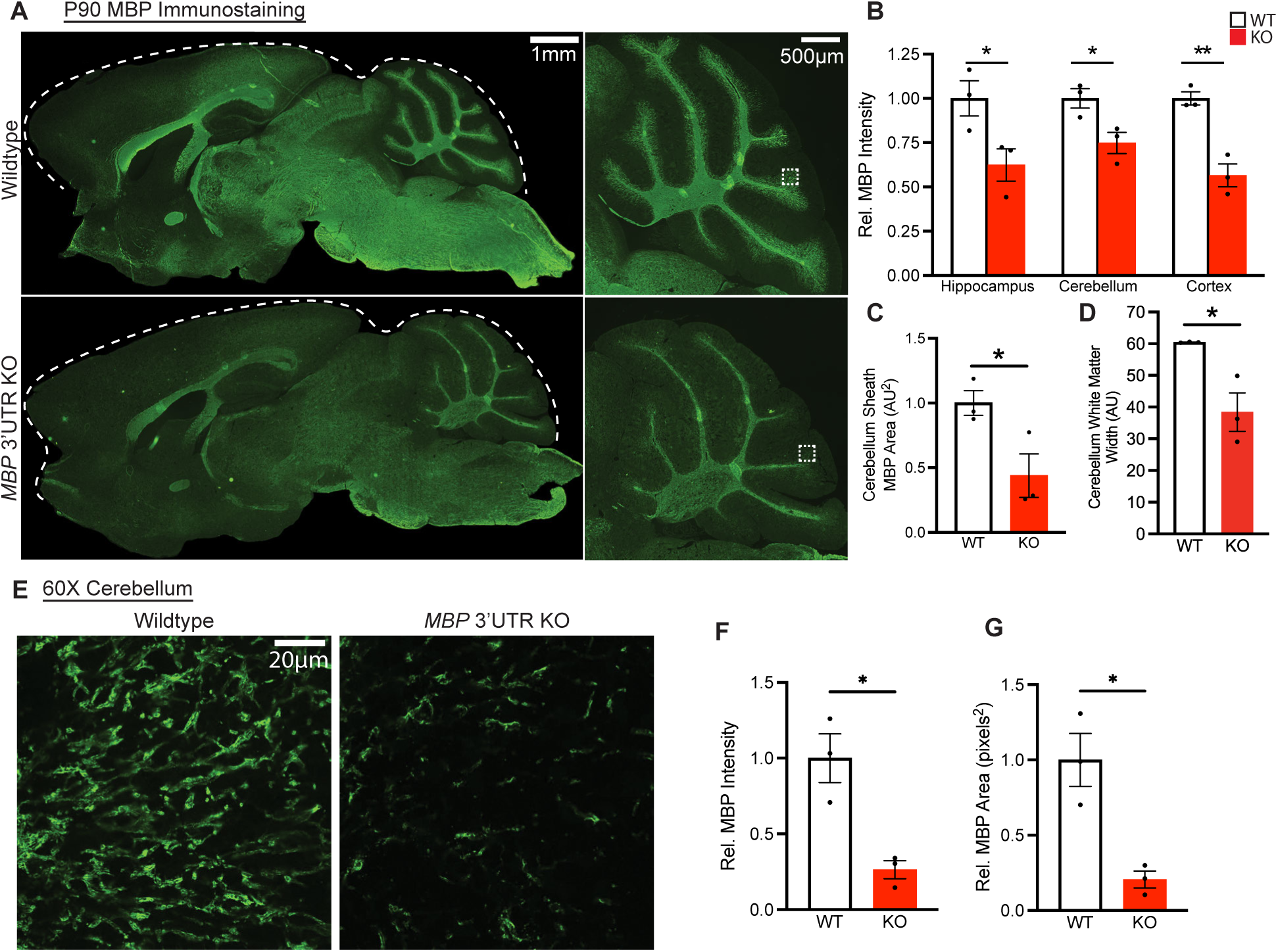
Immunofluorescence reveals that *Mbp* 3’ UTR KOs are hypomyelinated. (A) MBP immunohistochemical staining of 3-month-old mouse brain imaged at 10X magnification. Dashed white line indicates brain boundary (n=3 mice per genotype). (B) Quantification of relative MBP fluorescence intensity in multiple brain regions (P=0.049, 0.036, 0.0042, individual Student’s t-tests). (C) MBP area in cerebellum sheath region (P=0.0089, Student’s t-test). (D) MBP white matter width in cerebellum (P=0.022, Student’s t-test). (E) Images of MBP immunohistochemical staining from P90 mouse cerebellum. High magnification (60X). (F) MBP intensity quantification in hi-mag cerebellum image (P=0.013, Student’s t-test). (G) MBP area quantification in hi-mag cerebellum image (P=0.013, Student’s t-test). Data represents mean ± SEM. *P < 0.05, **P < 0.01, ***P < 0.001, ****P< 0.0001.

Next, to examine myelin ultrastructure in greater detail, we performed transmission electron microscopy (TEM) on cross sections of optic nerves from 3-month-old mice (Fig. 6A). Myelin thickness can be quantified using the g-ratio, which is calculated from the axon diameter divided by the total fiber (axon and myelin sheath) diameter. Thus, in the absence of axon diameter differences, a smaller g-ratio corresponds to thicker myelin. To robustly and accurately analyze g-ratio, we implemented a custom Matlab analysis pipeline (40) that corrects for non-circular myelin cross-sections by calculating all possible radii of a given sheath from the inner and outer myelin diameter coordinates. Using this program, we found that, compared to wildtype mice, homozygous *Mbp* 3’ UTR KO mice have a significantly higher average g-ratio (Fig. 6B and 6C). Because no difference was observed in axon diameter (Fig. 6D), this higher g-ratio indicates that homozygous *Mbp* 3’ UTR KO mice have thinner myelin sheaths. In addition, the number of unmyelinated axons was significantly increased in homozygous *Mbp* 3’ UTR KO mice (Fig. 6E).

**Figure 6.**
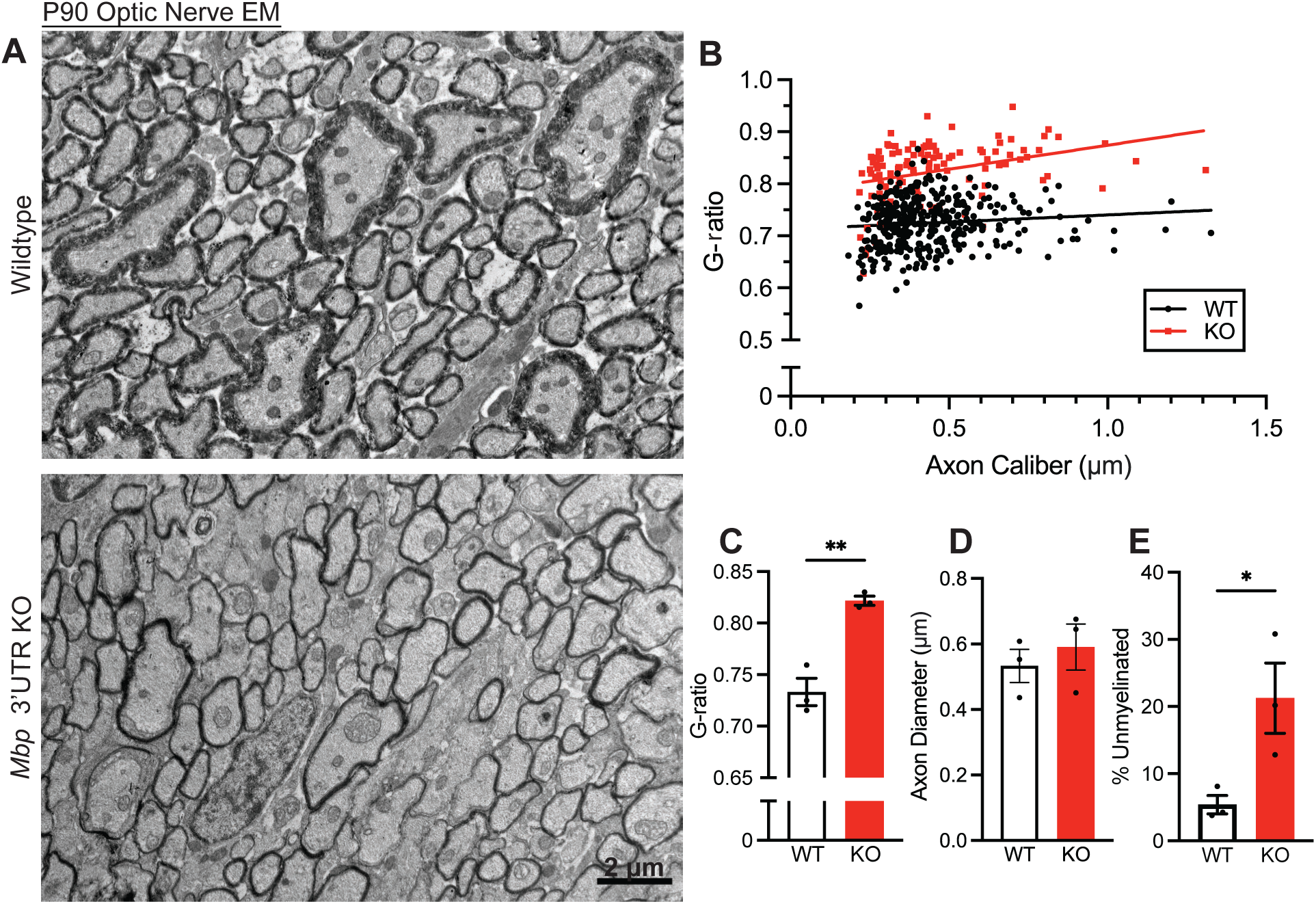
EM reveals that *Mbp* 3’ UTR KOs are hypomyelinated. (A) 3-month-old mouse optic nerve EMs (2000x). (B) Scatterplot of axon caliber vs. G-ratio. Each data point represents one myelinated axon (n=3 mice per genotype, 37–77 axons per image). (C) Optic nerve G-ratio (P=0.0032, Student’s t-test). (D) Optic nerve axon diameters (P=0.54, Student’s t-test). (E) Percent of unmyelinated axons (P=0.043, Student’s t-test). Data represents mean ± SEM. *P < 0.05, **P < 0.01, ***P < 0.001, ****P< 0.0001.

To determine the effect of *Mbp* 3’ UTR replacement on the peripheral nervous system (PNS), we also performed TEM on 3-month-old sciatic nerves (Fig. S5A). G-ratio analyses revealed no difference between wildtype and homozygous *Mbp* 3’ UTR KO mice (Fig. S5B and Fig. S5C). We also observed no difference in axon diameter (Fig. S5D), which indicates that myelin thickness did not change. In addition, the percentage of unmyelinated axons remained low with no significant differences between wildtype and homozygous *Mbp* 3’ UTR KOs (Fig. S5E). These results are consistent with the lack of PNS phenotypes in *shiverer* mice (12, 41), which has been attributed to compensation by the Schwann-cell specific myelin protein, myelin protein zero (MPZ, also known as protein zero or P0), which also functions in myelin compaction in the PNS (42).

### *Mbp* 3’ UTR KO mice exhibit motor and learning defects

After observing that homozygous *Mbp* 3’ UTR KO mice have tremors (Fig. 3F and 3G), defects in *Mbp* mRNA transport and local translation (Fig. 4), and hypomyelination across multiple brain regions (Fig. 5), we asked whether replacing the *Mbp* 3’UTR results in other functional and behavioral consequences *in vivo*. First, to assess motor coordination and balance, we tested wildtype and homozygous *Mbp* 3’ UTR KO mice on the Rotarod over two days of training with variable speed and a third trial day with constant speed. We found that homozygous *Mbp* 3’ UTR KO mice, compared to wildtype mice, quickly fall off the Rotarod (Fig. 7A). These results suggest that *Mbp* 3’ UTR KO mice have functional deficits in motor coordination and balance.

**Figure 7.**
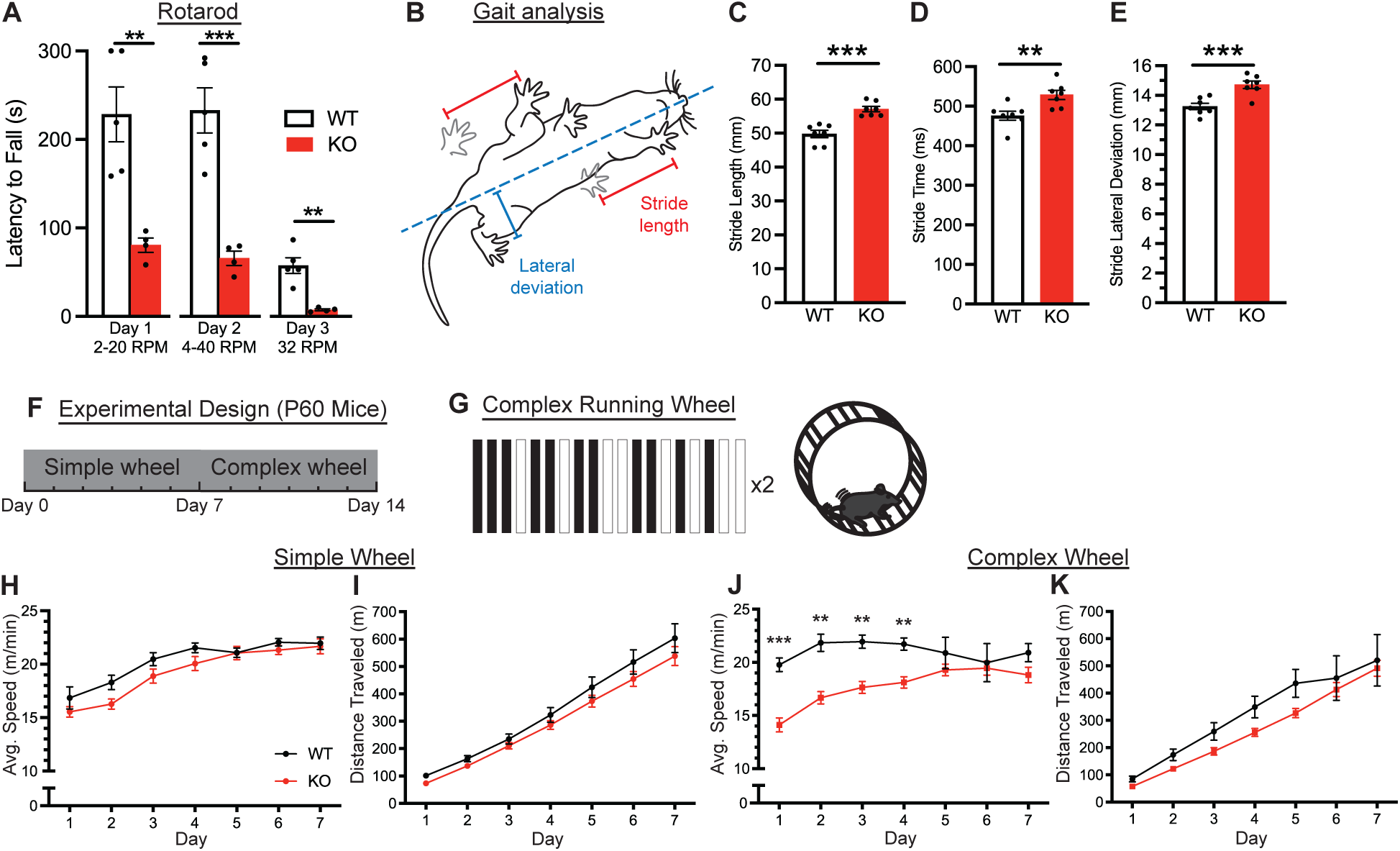
*Mbp* 3’ UTR KO mice display impaired coordination and motor learning. (A) Rotarod latency to fall from 3-months-old mice trained on day 1 and 2 at increasing speed. Day 3 test is at constant speed (P=0.0045, 0.0008, 0.0017, individual Student’s t-tests). (B) Schematic of gait analysis metrics. Stride length (indicated in red) is the distance between the previous step (gray paw prints) and the following step. Lateral deviation (indicated in blue) is the distance of the inside of the paw from the midline (dashed blue line, from nose to tail). (C) Stride length (P=0.0002, Student’s t-test, n=8 animals per genotype). (D) Stride time (P=0.007, Student’s t-test, n=8 animals per genotype). (E) Stride lateral deviation (P=0.0009, Student’s t-test, n=8 animals per genotype). (F) Experimental design. P60 wildtype and *Mbp* 3’ UTR KO mice were single housed and introduced to the simple running wheel for 1 week. The following week, they were introduced to the complex wheel for 1 week (n=8 mice per genotype). (G) Schematic of the complex wheel pattern. The pattern repeats twice in one revolution of the wheel. (H) Average running speed of wildtype and *Mbp* 3’ UTR KO mice on a simple wheel for 7 days (Two-way ANOVA with Šídák’s multiple comparisons test). (I) Average running speed of wildtype and *Mbp* 3’ UTR KO mice on a complex wheel for 7 days (Two-way ANOVA with Šídák’s multiple comparisons test). (J) Cumulative running distance of wildtype and *Mbp* 3’ UTR KO mice on a simple wheel for 7 days (Two-way ANOVA with Šídák’s multiple comparisons test). (K) Cumulative running distance of wildtype and *Mbp* 3’ UTR KO mice on a complex wheel for 7 days (Two-way ANOVA with Šídák’s multiple comparisons test). Data represents mean ± SEM. *P < 0.05, **P < 0.01, ***P < 0.001, ****P< 0.0001.

Next, we further assayed these motor defects by analyzing the gait of homozygous *Mbp* 3’ UTR KO mice (Fig. 7B). Compared to wildtype mice, they exhibited a longer and wider stride, as well as took longer to make each step (Fig. 7C–7E). These functional differences may help homozygous *Mbp* 3’ UTR KO mice to continue ambulating despite persistent tremors and hypomyelination.

To test for any changes in voluntary locomotion, we performed the open field assay (Fig. S6A and S6B). Over a 2-hour period, we observed no statistically significant differences in the distance traveled between wildtype, heterozygous, and homozygous *Mbp* 3’UTR KO mice (Fig. S6C, S6D), indicating no differences in voluntary locomotor activity. In addition, we also observed no differences in the time spent in the center (Fig. S6E–S6H), which is consistent with a lack of thigmotaxis (i.e., staying near walls), which can be a sign of anxiety-like behavior. Thus, open field results indicate that the absence of the *Mbp* 3’ UTR does not produce behavioral changes in voluntary locomotion or anxiety-like behavior.

Finally, to investigate whether replacing the *Mbp* 3’ UTR affects motor learning, we employed a complex wheel running task (43) (Fig. 7F and 7G), which is dependent on adaptive myelination (44). Beginning at P60, for 7 days, wildtype and homozygous *Mbp* 3’ UTR KO animals were housed in a cage containing a standard, simple running wheel in which all rungs of the wheel are evenly spaced. Across one week of training on the simple wheel, wildtype and homozygous *Mbp* 3’ UTR KO mice ran with no significant differences in average speed (Fig. 7H) or distance (Fig. 7I). Thus, despite persistent tremors and an altered gait, homozygous *Mbp* 3’ UTR KO mice are capable of similar running behavior on simple wheels. After one week, the simple wheel was replaced with a complex running wheel, which has rungs removed in an asymmetric pattern (Fig. 7G). On the complex wheels, homozygous *Mbp* 3’ UTR KO mice ran similar overall distances as wildtype mice. However, they exhibited significantly lower average speeds for the first four days of testing (Fig. 7J and 7K), which strikingly indicates a delay in motor learning in the absence of the *Mbp* 3’ UTR.

## DISCUSSION

Here, to test the hypothesis that the *Mbp* 3’ UTR is required for *in vivo* myelination in a mammalian system, we generated a transgenic mouse in which the endogenous *Mbp* 3’ UTR was replaced with stabilizing SV40 polyA sequences (Fig. 3A). Notably, homozygous *Mbp* 3’ UTR KO mice display persistent tremors (Fig. 3I and 3J), prompting further investigation into the cellular and functional consequences of 3’ UTR loss. Homozygous *Mbp* 3’ UTR KO mice are hypomyelinated by multiple metrics and oligodendrocytes cultured from these mice display intrinsic cellular defects. Immunostaining against MBP revealed visibly lower expression of this protein in multiple brain regions in homozygous *Mbp* 3’ UTR KO mice compared to wildtype (Fig. 4), indicative of widespread hypomyelination. Electron microscopy showed reduced myelin thickness and a greater proportion of unmyelinated axons in the optic nerve of homozygous *Mbp* 3’ UTR KO mice compared to wildtype (Fig. 5). These *in vivo* observations are consistent with defects in *Mbp* mRNA transport and translation exhibited in 3D cultured primary oligodendrocytes from homozygous *Mbp* 3’ UTR KO mice (Fig. 6A–E). Together, these results indicate that the 3’ UTR is critical for developmentally stereotyped *Mbp* mRNA localization and translation in oligodendrocytes, and that its loss leads to cell-autonomous myelination defects that extend to hypomyelination across multiple brain regions.

### Effects of 3’ UTR Replacement on Transcript Levels

Unexpectedly, *Mbp* CDS expression levels in homozygous *Mbp* 3’ UTR KO mice were reduced to ∼40% of wildtype levels (Fig. 3D and 3E). Thus, one potential concern about our *Mbp* 3’ UTR KO model is that the observed phenotypes could simply reflect partial loss of MBP function (a hypomorphic effect), rather than a specific defect in mRNA localization. However, this is contradicted by the lack of phenotypes in several *Mbp* rodent models. First, a recent study generated 6 mouse lines with various levels of *Mbp* mRNA reduction (ranging from 93–11 %) by excising regions in the *Mbp* enhancer but leaving the 3’ UTR intact (35). Of these, the line expressing 51% *Mbp* mRNA, the closest level to our homozygous *Mbp* 3’ UTR KO mice, displayed no significant changes in g ratio and no behavioral phenotypes. Another line with 22% *Mbp* mRNA also had no behavioral phenotypes. The line with the lowest *Mbp* mRNA level (11%) displayed tremors around P18. Second, shiverer mice have been extensively studied for decades. Heterozygous *shiverer* mice express lower levels of *Mbp* mRNA than wildtype mice (36). Behaviorally, heterozygous *shiverer* mice perform better on the Rotarod than homozygous *shiverer* mice and improve over days of training (Kuhn et al., 1995), whereas *Mbp* 3’ UTR KO mice do not improve with Rotarod training (Fig. 7A). Third, the *mld* (myelin deficient) mouse line also harbors *Mbp* mutations resulting in reduced *Mbp* mRNA levels, but these mice also have a comparable myelin composition to wildtype controls with no reported behavioral phenotypes (45). The absence of haploinsufficiency in these *Mbp* hypomorphs is perhaps consistent with the extraordinarily high wildtype levels of *Mbp* mRNA, which is the highest expressed mRNA in oligodendrocytes and is ∼10-fold higher than the second most abundant mRNA by bulk RNA-seq (45, 46). Thus, the lack of tremors and behavioral phenotypes in multiple *Mbp* mutant lines with reduced *Mbp* mRNA levels supports our conclusion that phenotypes in our homozygous *Mbp* 3’ UTR KO mice can be attributed to defects in mRNA localization.

Decreased levels of *Mbp* CDS mRNA in homozygous *Mbp* 3’ UTR KO mice may also suggest that the 3’ UTR enhances the stability of *Mbp* mRNA. Our data is consistent with a study in zebrafish comparing the expression of a reporter construct containing the *Mbp* 3’ UTR versus one containing the SV40 polyA sequence; the SV40 polyA construct was found to have lower expression in both the cell body and the myelin sheath. Moreover, comparing to other myelin-enriched 3’ UTRs, SV40 polyA construct expression was significantly lower (Yergert et al., 2021). Thus, myelin-specific 3’ UTRs may promote higher mRNA abundance especially in native cellular environments (e.g., the oligodendrocyte), in part through effects on transcript stability, potentially exceeding that conferred by SV40 polyA. Importantly, replacement of an endogenous 3’UTR with SV40 polyA does not universally result in reduced CDS expression. For example, 3’UTR replacement of *importin β1* and *mTOR* – genes whose localization and translation are regulated by their 3’UTRs in response to axonal injury – did not alter CDS mRNA levels (32, 33). Together, these findings indicate that the impact of SV40 polyA replacement on mRNA abundance may be gene- and context-dependent, and suggest that the Mbp 3’UTR has a uniquely important role in maintaining physiological Mbp mRNA levels in oligodendrocytes.

### Dissecting Elements within the *Mbp* 3’ UTR

Replacement of the large ∼1.5-kb 3’ UTR of mouse *Mbp* mRNA likely removed multiple regions responsible for mRNA stability, localization, and modulation of translation. Early experiments identified two distinct regions that affect mRNA transport – the RTS (RNA transport signal), located ∼400bp into the 3’ UTR and the RLR (RNA localization region), located ∼1100bp into the 3’ UTR (31). Additional motifs for RNA-binding proteins, such as hnRNP E1 and hnRNP K have also been identified within the *Mbp* 3’ UTR (47). A recent study that systematically tested many Mbp 3’UTR variants in parallel combined with DMS-MaPseq to solve the RNA structure of *Mbp* 3’ UTR revisited the question of what elements within the 3’ UTR are important for *Mbp* mRNA transport. This study did not replicate the necessity of the RTS for *Mbp* mRNA transport, but it did identify a ∼120-bp structure named the MLS (Mbp localization sequence) that partially overlaps with the RTS as being necessary and sufficient for *Mbp* mRNA localization (34). Surprisingly, this region associates with hnRNP F, which regulates translation initiation, and its association with *Mbp* mRNA was hypothesized to switch the RNA from a transport complex to a translation complex (34). Thus, validation of these motifs and regions as well as their associated partner RNA-binding proteins could further dissect their contributions to the distinct processes of RNA stabilization, transport, and translation. These insights could lead to the refinement of a mouse model targeting smaller deletion regions within the 3’ UTR.

### MBP Local Translation and Motor Learning

Our study shows that homozygous *Mbp* 3′ UTR KO mice display deficits in motor learning on the complex wheel assay, a task that has been linked to adaptive myelination during skill acquisition (44). This finding implicates MBP local translation not only in myelin maintenance, but also in experience- and activity-dependent myelin remodeling. Consistent with a role for MBP local translation in myelin maintenance, recent work showed that MBP expression is critical for continuous myelin synthesis at the inner tongue and for long-term sheath stability (48).

At the cellular level, neuronal activity has been shown to engage signaling pathways that regulate MBP local translation. Specifically, membrane-associated Fyn kinase is activated by neuronal activity and promotes MBP translation in oligodendrocytes (49). Mechanistically, Fyn-mediated phosphorylation of hnRNP F causes its dissociation from Mbp mRNA, thereby relieving translational repression (50). Together with the identification of the association of hnRNP F with the MLS of *Mbp* 3’ UTR (34), these studies collectively support a model in which *Mbp* mRNAs transported to the myelin sheath are locally poised to respond to neuronal activity-induced kinase activation by initiating translation.

This link between motor learning and MBP local translation is also consistent with *in vivo* data across many motor learning experiments. First, rats trained for 11 days at a skilled reaching task had elevated MBP staining near the motor cortex. Higher MBP staining also correlated with faster learning rates (51). Second, oligodendrocytes generated after complex wheel training express *Mbp* mRNA along nascent myelin sheaths (52). Mice trained on the complex wheel also have elevated *Mbp* mRNA levels in their motor cortex and higher *Mbp* mRNA levels correlate with higher training success rates (53). Thus, both cellular and systems-level studies are consistent with a role for MBP local translation in response to motor learning and neuronal activity.

Holistically, our work provides an important link between the cellular role of *Mbp* 3’ UTR in mRNA localization and local translation and its *in vivo* role in myelination and motor learning. As we learn more about 3’ UTR structure and regulation, future work can yield important mechanistic insights connecting molecular mechanisms with whole animal behavior.

## ONLINE METHODS

### Generation of transgenic mice

Mice were created by generating guide RNAs for the region flanking the 3’ UTR of the *Mbp* gene. A donor construct consisting of four SV40 PolyA sequences flanked by regions homologous to the regions flanking the *Mbp* 3’ UTR was produced. A transgenic mouse was generated after homology-directed repair (HDR) or homologous recombination. Though this is technically a knock-in mouse, its effect is to disrupt the function of the 3’ UTR and thus we call it the *Mbp* 3’ UTR “knockout” (KO) mouse.

### Mouse genotyping and DNA gels

Toe clips were collected from mice between P5-P7. DNA was extracted and amplified using REDExtract-N-Amp PCR ReadyMix (Sigma R4775) and run on a 3% agarose gel containing ethidium bromide. The following primers were used: MBP WT Fwd (5’ CTCTTAATCCCGTGGAGCCG 3’), MBP WT Rev (5’ CAGCCTGTGCTCACATACCA 3’), MBP KO Fwd (5’ CTCCCCGCGTTGTTAACTTGTT 3’), and MBP KO Rev (5’ GTGCTTCAGGATCCAGACATGAT 3’).

### qPCR

Mouse brains were dissected and mRNA was isolated using RNeasy Kit (QIAGEN). mRNA was measured using Quant-iT RNA Assay Kit (Thermo Q33225) on a fluorescence plate reader and converted to cDNA using the Superscript IV VILO kit (Thermo 11756050). All reactions were run on QuantStudio 3 (Thermo) using Powerup Sybr Green (Thermo A25778).

All primers were validated by testing their efficiencies. The following primers against *Mbp* were used: MBP_Exon1_Fwd #3 (5’ TCACAGAAGAGACCCTCACA 3’) and MBP_Exon1_Rev #3 (5’ CCCTGTCACCGCTAAAGAAG 3’). Results from the qPCR were normalized against control reactions against GAPDH and HGPRT. The following control primers were used: GAPDH mouse Fwd (5’ TCTCTGCTCCTCCCTGTTCC 3’), GAPDH mouse Rev (5’ GATGGTGATGGGCTTCCCGT 3’), HGPRT Fwd (5’ TCAGTCAACGGGGGACATAAA 3’), and HGPRT Rev (5’ GGGGCTGTACTGCTTAACCAG 3’).

### Mouse brain immunohistochemistry

Mice were anesthetized with a cocktail of ketamine (100 mg/kg) and xylazine (20 mg/kg) then transcardially perfused at a rate of 3mL/min, first with 1X phosphate-buffered saline followed by 4% paraformaldehyde (PFA, Electron Microscopy Services, 15711). Brains were removed and placed in 4% PFA overnight at 4°C, then washed with 1X PBS and transferred to 30% sucrose. Frozen tissue was mounted in Tissue Plus O.C.T. Compound (Fisher HealthCare, 23-730-571) and frozen at −80°C until sectioning. Blocks were sectioned to 12-μm thick slices, fixed in 4% PFA for 10 minutes at room temperature, and blocked with 10% donkey serum in 0.1% Triton X-100 in PBS for 30min. Sections were incubated at 4°C overnight with the following primary antibodies at these concentrations: MBP (Abcam ab7349, 1:100), PLP (Novus NBP1-87781, 1:250, CNP (Sigma C5922, 1:250). After, the sections were incubated in secondary antibodies (Jackson ImmunoResearch Laboratories, 1:200) for 2 hours at room temperature. A DAPI stain (Sigma D9542) was applied for 3 minutes at room temperature. Sections were mounted in Vectashield Plus Mounting Medium (Vector Laboratories, H-1900).

### Immunopanning and culturing of primary oligodendrocytes

Oligodendrocyte precursor cells (OPCs) were purified from Sprague-Dawley rat pups (P6–P8) by immunopanning as previously described in Dugas and Emery (2013). Briefly, cortical tissue was dissociated by papain digestion then filtered through a Nitex mesh to obtain a mixed single-cell suspension. This suspension was incubated in 2 negative-selection plates coated with anti-Ran-2 and anti-GC antibodies, then in a positive-selection plate coated with anti-O4 antibody. Adherent cells were trypsinized and cultured in proliferation media containing PDGF and NT-3 or differentiation media containing T3 (triiodothyrone, a derivative of thyroid hormone). For 3D cultures, 2-mm microfiber inserts for 12-well plates were washed in ethanol, then coated in poly-D-lysine (PDL) hydrobromide, then cultured for 14 days at 37°C.

### Oligodendrocyte smFISH & immunofluorescence staining

Oligodendrocytes were fixed on glass coverslips in 4% paraformaldehyde, 4% sucrose, 1X PHEM (60mM PIPES, 25mM HEPES, 10mM EGTA, 2mM MgSO4) for 10 minutes at 37°C. Cells were permeabilized in 70% ethanol at 4°C for 1 hour. Coverslips were incubated with probes against Mbp mRNA (Stellaris, 1:100) and primary antibodies for Tubulin (1:1000 Sigma, T9026-2mL), and MBP (1:100 Abcam, ab7349) at 37°C for 4–16 hours in Hybridization buffer (Dextran sulfate, SSC buffer, deionized formamide). Coverslips were incubated with secondary antibodies (Jackson ImmunoResearch Laboratories, 1:200) and Phalloidin dye (Alexa Fluor Plus 647 Phalloidin, Thermo Fisher A30107, 1:400) for 30 minutes. Coverslips were mounted on glass slides with GLOX Anti-Fade (glucose, Tris-HCl, SSC buffer, catalase, glucose oxidase) for imaging. Micrographs were acquired on a Nikon Ti2 spinning-disk confocal using 20X, 60X, or 100X objectives with a scMOS Hamamatsu ORCA-FusionBT camera.

### Colocalization analysis

Confocal images of DIV 3–6 oligodendrocytes were analyzed by counting *Mbp* mRNA granules in a bounding box. First, RNA puncta that were colocalized with tubulin were counted. Then, remaining RNA puncta that were colocalized with phalloidin staining for actin were counted.

### Electron microscopy

In brief, mice were transcardially perfused with 2% glutaraldehyde + 2% paraformaldehyde in 0.1M cacodylate buffer at pH 7.4. Optic and sciatic nerves were dissected out, incubated in fixative overnight at 4°C. Tissue was sectioned into 100 µm slices and imaged with a JEOL 1200 EXII Transmission Electron Microscope at 250X and 1000X.

### Tremor analysis

1-minute videos were taken of 2-month-old mice in a walled enclosure (13”L × 10.5”W x 3”H). An investigator blinded to genotype was asked to score each video using a stopwatch to record total time tremoring.

### Rotarod running wheel

Mice were tested 3 times per day on a Rotarod (Ugo Basile, Milan, Italy) for 3 consecutive days with a minimum of 5 minutes rest between each trial. Day 1 training consisted of 1 minute of 2 rpm, followed by acceleration to 20 rpm over 300 seconds. Day 2 testing consisted of an acceleration from 4–40 rpm over 300 seconds. Day 3 consisted of a test at a constant speed of 32 rpm over 300 seconds. An average latency to fall was taken from each trial day for each genotype.

### Gait analysis

Each mouse was placed into an enclosed treadmill apparatus (CSI-G-TRD, Columbus Instruments and CleverSys) with a camera to record an underside view. Training consisted of ramping the speed up to 8 cm/s and walking for 15 seconds. After 1 minute of rest, the treadmill speed was ramped up to 8 cm/s and a 20 second recording was started. Apparatus was cleaned with 70% ethanol between trials. Gait was analyzed using Treadscan software (CleverSys).

### Open field assay

At the beginning of each trial, a single mouse was placed in a clear plexiglass container (custom, NIMH Behavioral Core) and recorded from above for the duration of a 2-hour trial. Videos were analyzed using TopScan 3.0 (CleverSys).

### Complex wheel assay

Cages containing a running wheel and digital recording devices (Lafayette Neuroscience) were purchased to monitor running speed of mice. Complex wheels were assembled as previously described (McKenzie et al. 2014), briefly: 16 rungs were removed from a 38-rung, 12.7cm diameter running wheel to create a 19-rung pattern that repeats twice per revolution of the running wheel. Mice were maintained on a 12h/12h light/dark cycle with food and water provided ad-libitum. Food and water were monitored every 1–2 days during the “lights on” period of the cycle so as not to disturb running.

### Statistical analysis

Means, standard deviations, standard errors, and statistical analyses were calculated and graphed using Prism 9 (GraphPad). Data were first tested for normality or parametricity using Shapiro-Wilk and Kolmogorov-Smirnov tests. Normal data was analyzed using t-test or one-way ANOVA. Non-normal data was analyzed using the Kruskal-Wallis test or Mann-Whitney test.

## AUTHOR CONTRIBUTIONS

Conceptualization – M.-m.F.;

Methodology – J.C.N., L.M., M.-m.F.;

Formal analysis – J.C.N., L.M., H.B., H.S.R., H.N., M.-m.F.;

Investigation – J.C.N., L.M., A.V. H.N., M.-m.F;

Writing – Original Draft, J.C.N., M.-m. F.;

Writing – Review & Editing, J.C.N., H.B., M.-m. F.;

Visualization – J.C.N., H.B., L.M., H.N., M.-m.F.;

Supervision – M.-m.F.;

Funding Acquisition – M.-m.F.

## DECLARATION OF INTERESTS

The authors declare no competing interests.

## ACKNOWLEDGEMENTS

We thank the NIMH (National Institute of Mental Health) Rodent Behavior Core (Yogita Chudasama, Sean Bradley) for training and insightful discussions. We thank the NINDS (National Institute of Neurological Disorders and Stroke) Electron Microscopy Core (Susan Cheng, Sandra Moreira) for training and assistance with TEM. We thank the following NINDS labs for use of their equipment/reagents, technical training, and mentorship: Antonina Roll-Mecak Lab, Kenton Swartz Lab (Helena Tsg-Hui Chang), Joseph Mindell Lab, John Hammer Lab (Sri Repudi). We thank the Porter Neuroscience Research Center Animal Core Facility (Heather Narver, Maria Lorenzo, Stephanie Levin, Annie Xin Wu) for animal care and assistance with animal transfers.

M.M.F. is supported by NINDS intramural funding (NS009432) and the Human Frontier Science Program (RGEC32/2023). The NIMH Rodent Behavioral Core is supported by MH002952.

## Notes

### Competing Interest Statement

The authors have declared no competing interest.

## REFERENCES

1. K.-A. Nave, H. B. Werner, Myelination of the Nervous System: Mechanisms and Functions. Annu. Rev. Cell Dev. Biol. 30, 503–533 (2014).

2. A. Peters, The formation and structure of myelin sheaths in the central nervous system. J. Biophys. Biochem. Cytol. 8, 431–446 (1960).

3. N. Baumann, D. Pham-Dinh, Biology of Oligodendrocyte and Myelin in the Mammalian Central Nervous System. Physiol. Rev. 81, 871–927 (2001).

4. S. Aggarwal, et al., Myelin Membrane Assembly Is Driven by a Phase Transition of Myelin Basic Proteins Into a Cohesive Protein Meshwork. PLOS Biol. 11, e1001577 (2013).

5. E. H. Eylar, S. Brostoff, G. Hashim, J. Caccam, P. Burnett, Basic A1 Protein of the Myelin Membrane: THE COMPLETE AMINO ACID SEQUENCE. J. Biol. Chem. 246, 5770–5784 (1971).

6. O. Jahn, et al., The CNS Myelin Proteome: Deep Profile and Persistence After Post-mortem Delay. Front. Cell. Neurosci. 14 (2020).

7. A. Roach, N. Takahashi, D. Pravtcheva, F. Ruddle, L. Hood, Chromosomal mapping of mouse myelin basic protein gene and structure and transcription of the partially deleted gene in shiverer mutant mice. Cell 42, 149–155 (1985).

8. S. M. Molineaux, H. Engh, F. de Ferra, L. Hudson, R. A. Lazzarini, Recombination within the myelin basic protein gene created the dysmyelinating shiverer mouse mutation. Proc. Natl. Acad. Sci. U. S. A. 83, 7542–7546 (1986).

9. E. Barbarese, M. L. Nielson, J. H. Carson, The Effect of the Shiverer Mutation on Myelin Basic Protein Expression in Homozygous and Heterozygous Mouse Brain. J. Neurochem. 40, 1680–1686 (1983).

10. F. Biddle, E. March, J. R. Miller, Research News. Mouse News Lett. 48, 24 (1973).

11. A. Privat, C. Jacque, J. M. Bourre, P. Dupouey, N. Baumann, Absence of the major dense line in myelin of the mutant mouse ‘shiverer.’ Neurosci. Lett. 12, 107–112 (1979).

12. J. Rosenbluth, Central myelin in the mouse mutant shiverer. J. Comp. Neurol. 194, 639–648 (1980).

13. C. Readhead, et al., Role of Myelin Basic Protein in the Formation of Central Nervous System Myelin. Ann. N. Y. Acad. Sci. 605, 280–285 (1990).

14. G. F. Chernoff, Shiverer: an autosomal recessive mutant mouse with myelin deficiency. J. Hered. 72, 128–128 (1981).

15. P. l. Kuhn, E. Petroulakis, G. a. Zazanis, R. d. McKinnon, Motor function analysis of myelin mutant mice using a rotarod. Int. J. Dev. Neurosci. 13, 715–722 (1995).

16. H. D. Shine, C. Readhead, B. Popko, L. Hood, R. L. Sidman, Morphometric analysis of normal, mutant, and transgenic CNS: correlation of myelin basic protein expression to myelinogenesis. J. Neurochem. 58, 342–349 (1992).

17. S.-K. Song, et al., Dysmyelination Revealed through MRI as Increased Radial (but Unchanged Axial) Diffusion of Water. NeuroImage 17, 1429–1436 (2002).

18. D. R. Colman, G. Kreibich, A. B. Frey, D. D. Sabatini, Synthesis and incorporation of myelin polypeptides into CNS myelin. J. Cell Biol. 95, 598–608 (1982).

19. K. Ainger, et al., Transport and localization of exogenous myelin basic protein mRNA microinjected into oligodendrocytes. J. Cell Biol. 123, 431–441 (1993).

20. J. H. Carson, K. Worboys, K. Ainger, E. Barbarese, Translocation of myelin basic protein mRNA in oligodendrocytes requires microtubules and kinesin. Cell Motil. 38, 318–328 (1997).

21. L. M. Meservey, V. V. Topkar, M. Fu, mRNA Transport and Local Translation in Glia. Trends Cell Biol. 31, 419–423 (2021).

22. N. Snaidero, et al., Myelin Membrane Wrapping of CNS Axons by PI(3,4,5)P3-Dependent Polarized Growth at the Inner Tongue. Cell 156, 277–290 (2014).

23. A. Valenzuela, L. Meservey, H. Nguyen, M. Fu, Golgi Outposts Nucleate Microtubules in Cells with Specialized Shapes. Trends Cell Biol. 30, 792–804 (2020).

24. M. Weigel, L. Wang, M. Fu, Microtubule organization and dynamics in oligodendrocytes, astrocytes, and microglia. Dev. Neurobiol. 81, 310–320 (2021).

25 . K. J. Chapple, et al., A myelinic channel system for motor-driven organelle transport. [Preprint] (2024). Available at: http://biorxiv.org/lookup/doi/10.1101/2024.06.02.591488 [Accessed 12 November 2025].

26. M. Fu, et al., The Golgi Outpost Protein TPPP Nucleates Microtubules and Is Critical for Myelination. Cell 179, 132–146.e14 (2019).

27. D. A. Lyons, S. G. Naylor, A. Scholze, W. S. Talbot, Kif1b is essential for mRNA localization in oligodendrocytes and development of myelinated axons. Nat. Genet. 41, 854–858 (2009).

28. A. L. Herbert, et al., Dynein/dynactin is necessary for anterograde transport of Mbp mRNA in oligodendrocytes and for myelination in vivo. Proc. Natl. Acad. Sci. 114, E9153–E9162 (2017).

29. M. Kimura, et al., Restoration of myelin formation by a single type of myelin basic protein in transgenic shiverer mice. Proc. Natl. Acad. Sci. U. S. A. 86, 5661–5665 (1989).

30. M. Kimura, et al., Overexpression of a minor component of myelin basic protein isoform (17.2 kDa) can restore myelinogenesis in transgenic *shiverer* mice. Brain Res. 785, 245–252 (1998).

31. K. Ainger, et al., Transport and Localization Elements in Myelin Basic Protein mRNA. J. Cell Biol. 138, 1077–1087 (1997).

32. R. B.-T. Perry, et al., Subcellular Knockout of Importin β1 Perturbs Axonal Retrograde Signaling. Neuron 75, 294–305 (2012).

33. M. Terenzio, et al., Locally translated mTOR controls axonal local translation in nerve injury. Science 359, 1416–1421 (2018).

34. V. V. Topkar, et al., RNA switch model for localization and translation of the myelin basic protein mRNA. [Preprint] (2025). Available at: http://biorxiv.org/lookup/doi/10.1101/2025.11.19.689361 [Accessed 19 December 2025].

35. H. Bagheri, et al., Myelin basic protein mRNA levels affect myelin sheath dimensions, architecture, plasticity, and density of resident glial cells. Glia 72, 1893–1914 (2024).

36. H. J. Roth, M. J. Hunkeler, A. T. Campagnoni, Expression of myelin basic protein genes in several dysmyelinating mouse mutants during early postnatal brain development. J. Neurochem. 45, 572–580 (1985).

37. M. E. Bechler, L. Byrne, C. ffrench-Constant, CNS Myelin Sheath Lengths Are an Intrinsic Property of Oligodendrocytes. Curr. Biol. 25, 2411–2416 (2015).

38. J.-M. Cioni, et al., Late Endosomes Act as mRNA Translation Platforms and Sustain Mitochondria in Axons. Cell 176, 56–72.e15 (2019).

39. Y.-C. Liao, et al., RNA Granules Hitchhike on Lysosomes for Long-Distance Transport, Using Annexin A11 as a Molecular Tether. Cell 179, 147–164.e20 (2019).

40. H. Richardson, HunRich/FindGRatio. (2023). Deposited 8 June 2023.

41. D. A. Kirschner, A. L. Ganser, Compact myelin exists in the absence of basic protein in the shiverer mutant mouse. Nature 283, 207–210 (1980).

42. Y. Ding, K. R. Brunden, The cytoplasmic domain of myelin glycoprotein P0 interacts with negatively charged phospholipid bilayers. J. Biol. Chem. 269, 10764–10770 (1994).

43. N. Hibbits, R. Pannu, T. John Wu, R. C. Armstrong, Cuprizone demyelination of the corpus callosum in mice correlates with altered social interaction and impaired bilateral sensorimotor coordination. ASN NEURO 1, e00013 (2009).

44. I. A. McKenzie, et al., Motor skill learning requires active central myelination. Science 346, 318–322 (2014).

45. J.-M. Roch, B. J. Cooper, R. Ramirez, J.-M. Matthieu, Expression of only one myelin basic protein allele in mouse is compatible with normal myelination. Mol. Brain Res. 3, 61–68 (1987).

46. Y. Zhang, et al., An RNA-Sequencing Transcriptome and Splicing Database of Glia, Neurons, and Vascular Cells of the Cerebral Cortex. J. Neurosci. 34, 11929–11947 (2014).

47. J. Torvund-Jensen, J. Steengaard, L. B. Askebjerg, K. Kjaer-Sorensen, L. S. Laursen, The 3’UTRs of Myelin Basic Protein mRNAs Regulate Transport, Local Translation and Sensitivity to Neuronal Activity in Zebrafish. Front. Mol. Neurosci. 11, 185 (2018).

48. M. Meschkat, et al., White matter integrity in mice requires continuous myelin synthesis at the inner tongue. Nat. Commun. 13, 1163 (2022).

49. H. Wake, P. R. Lee, R. D. Fields, Control of Local Protein Synthesis and Initial Events in Myelination by Action Potentials. Science 333, 1647–1651 (2011).

50. R. White, et al., Heterogeneous Nuclear Ribonucleoprotein (hnRNP) F Is a Novel Component of Oligodendroglial RNA Transport Granules Contributing to Regulation of Myelin Basic Protein (MBP) Synthesis. J. Biol. Chem. 287, 1742–1754 (2012).

51. C. Sampaio-Baptista, et al., Motor Skill Learning Induces Changes in White Matter Microstructure and Myelination. J. Neurosci. 33, 19499–19503 (2013).

52. L. Xiao, et al., Rapid production of new oligodendrocytes is required in the earliest stages of motor skill learning. Nat. Neurosci. 19, 1210–1217 (2016).

53. D. Kato, et al., Motor learning requires myelination to reduce asynchrony and spontaneity in neural activity. Glia 68, 193–210 (2020).

